# Engineering 3D neuronal networks with directional endogenous neuronal plasticity pathways

**DOI:** 10.1101/2023.05.17.540876

**Authors:** Gelson J Pagan-Diaz, Evin Kilacarslan, Matthew Wester, Saeedur Rahman, Onur Aydin, Lauren Gapinske, Yongdeok Kim, Daniel Buoros, M Saif A Taher, Rashid Bashir

## Abstract

The forward engineering of the structure and function of three-dimensional millimeter to centimeter scale living Neuronal Tissue Mimics (NTMs) can advance many engineering and biomedical applications. While hydrogels and 3D printing have achieved major breakthroughs in the development of cm-scale neural tissues that mimic structural morphologies in native neural networks, controlling and programming the resulting function of these NTMs have remained elusive. In this work, using human embryonic stem cell derived optogenetic neurons, we report the in-situ formation of the NTMs on a 2-dimensional micro electrode array with an intimate electrical contact between the electrodes and the tissue. These NTMs were optimized during the differentiation phase of the cells to enrich for neuronal populations that expressed receptors responsible for activating spike-timing dependent plasticity (STDP). Using an optical stimulation regiment with millisecond temporal and micrometer spatial resolution, we were able to program the otherwise omnidirectional spontaneous firing in the NTMs to demonstrate directional firing across different shapes of the NTMs. Our work can pave the way for developing cellular based computational devices, bio-processors, and biological memories.

## Introduction

In nature, “form drives function” is a consistent motif that has facilitated the discovery of many modalities of neuronal behavior, and has enabled the production of many neural-based models of the central nervous system^1^. For example, neuronal models have been developed capable of activating downstream tissues within engineered heterotypic multi-cellular engineered systems^2–5^. This is crucial, since the progress in tissue engineering depends on our ability to not only engineer isolated tissues that can mimic tissue-specific physiological behaviors but also their capability to interact with other systems as they do in native living systems. However, when these systems are constructed *in-vitro*, the pathways that would allow for “form” to “drive function” are disrupted. This results in tissue mimics that show the behaviors that the tissues would show in the mammalian system, but without the necessary organization in function that would allow them to function in unison with other interactive tissues. Furthermore, this is especially true for complex and highly dynamic tissues such as those formed from 3D neuronal networks. While these tissues have been shown to have high degrees of connectivity, synchrony, and bursting behaviors ^6–9^, they consistently show high variability in their resulting firing patterns. Furthermore, while Hebbian plasticity has been shown to be activated in *in-vitro* cultures to affect the dynamic of neuronal network at a cellular level^10^, achieving control of tissue-wide modulation of the directionality of neuronal activity has yet to be shown. This is also a crucial step towards development of large scale neuromorphic biohybrid systems^11^. The ability to enhance or control the electrophysiological functions of engineered neural circuits could be used for improvement in the emerging field of engineering biohybrid, neuronal-driven, biological machines. It would be highly advantageous to forward-engineer programmable neural networks that could be installed within *in vivo* or *in vitro* systems in order to achieve targeted functional behaviors ^12–14^.

To address the challenges mentioned above, here we show the viability of programming the direction of firing at large scale in millimeter-scale neural tissues by evoking spike-timing dependent plasticity (STDP). This form of Hebbian plasticity is induced by regulation of precise temporal correlations between action potentials of pre- and post-synaptic neuronal regions. Put simply, the Hebbian principle presents that when a presynaptic neuron fires consistently before its post synaptic target, this synapse undergoes long term potentiation (LTP), reinforcing the driving capacity of the presynaptic neuron unto the post-synaptic. Conversely, if a presynaptic neuron that consistently fires unto the post-synaptic after the post-synaptic has already been activated by an alternate presynaptic neuron, the aforementioned synapse will undergo long term depression (LTD) (**Supp. Figure 1a**). Furthermore, a well studied pathway of LTD induction involves strong glutamatergic activation of AMPA receptors, which causes a strong depolarization of the membrane which in turn causes the activation of NMDA receptors. NMDA receptor activation then causes a downstream activation cascade that results in the expression of more AMPA receptors facilitating communication across the synapse (**Supp. Figure 1b**). Our approach includes the recapitulation of these well-studied phenomena by using our recently reported biofabrication protocol of versatile 3D neuronal tissue mimics (NTMs), customization of electrode array layout, on-chip fabrication of NTMs, enrichment of neuronal populations that facilitated the eliciting of STDP, and high time-resolution optogenetic stimulation with sub-cellular spatial control. The effective combination of these aspects has allowed for a significant proof of concept towards forward-engineering 3-dimensional neuronal tissues whose large-scale firing patterns are programmable. Our work can have broad impact on developing biological neuromorphic engineered systems as memories or computational devices.

## Results and discussion

The implementation of optogenetic stimulation to evoke Hebbian plasticity was done with a Polygon1000 system (**Supp. Figure 2**; see Methods). This system can shine light across a 4 mm span capable of pixelating it down to ~10µm and scanning through them with millisecond resolution. With these capabilities, we stimulated beams of +50µm width with 465nm wavelength for 50 ms prior to stepping to the next section within <5ms intervals (**Figure 1a**). The minimum step width was selected as the thinnest beam that would not be affected (<5% thickness change) by refraction across the fluid, meaning that the actual width of the observed beam on the electrodes would be at least 95% of the designed value. On the other hand, the step timing was selected to coincide with the temporal parameters that have been reported to evoke long-term potentiation (LTP) through spike timing dependent plasticity (STDP).^15^ In summary, STDP-driven LTP results from a consistent activation of presynaptic neurons prior to activation of postsynaptic neurons^16^, and has been successfully and consistently shown in single neuron to neuron connections, primarily related to glutamatergic synaptic transmission^17^. A classic yet basic STDP induction protocol consists of stimulation at 100Hz without control of post-synaptic activity^18^. We implemented this protocol as a proof of concept to achieve STDP to bias the action potential firing direction (**Figure 1b**). We did so on compacted NTMs that had started spontaneously firing in random directions. Our goal was to potentiate synapses firing in the direction of the stepping stimulation, while synapses firing in opposite direction would be weakened through long-term depression (LTD) (**Figure 1c**).

**Figure 1.**
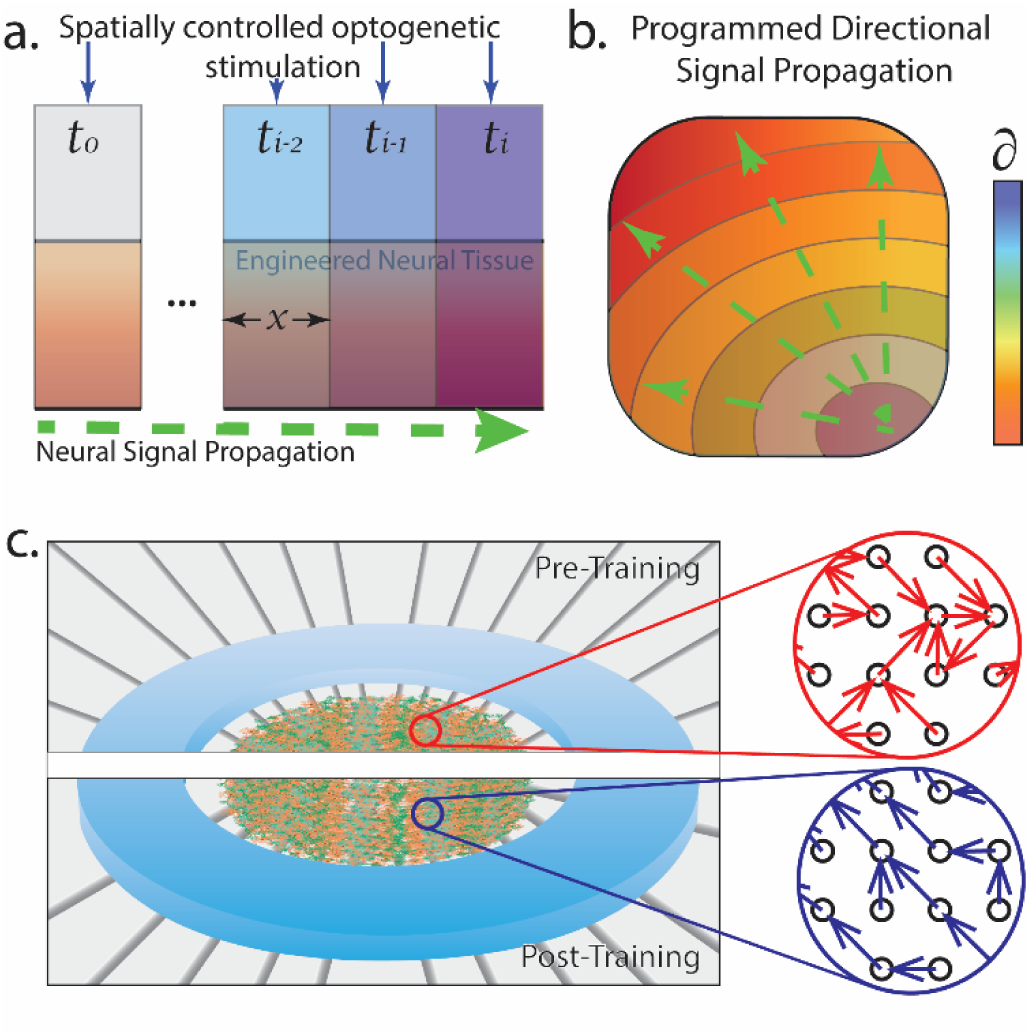
Using temporally and spatially controlled optogenetic stimulation to modulate the direction of action potential firing in engineering neural tissue mimics. **a.** Representative diagram showing proposed implementation of spatially and temporally optogenetic stimulation to evoke a directional firing pattern on NTMs (**b.**) where firing is effectively delayed by some δ interval. This would be implemented in an NTM with random firing pattern, and through guided stimulation synapses firing in opposing directions to the stimulation would be weakened, while the converse would be reinforced (**c**).

### Addition of Mouse Embryonic Fibroblasts (MEFs) to helps the compaction of NTMs enriched with glutamatergic neurons

The standard differentiation protocol used for NTM fabrication was enriched for motor neuronal identities (monitored by Hb9 expression)^19^. However, for the present approach, it was essential to enrich for glutamatergic neurons which could trigger STDP driven by activation of glutamatergic receptors.^18^ Few reports have successfully driven mESC differentiation towards near homogenous (~95%) of glutamatergic identity among neuronal populations^20–22^. Unfortunately, these studies employed an experimental design that finalized differentiation on a monolayer culture. Currently, our NTM fabrication protocol consists of differentiating ESCs as intact embryoid bodies prior to dissociation (done right after the beginning neural specialization)^23^. Dissociated differentiated neurons are then mixed with a liquid protein solution that, when incubated, cross-links as a hydrogel that provides a scaffold for neurons to undergo synaptogenesis and network maturation (**Fig 2a**). Therefore, our first goal was to optimize our protocol to achieve a high percentage of glutamatergic neural populations that would still compact under our biofabrication protocols (**Supp. Figure 3**). The primary alteration to the differentiation protocol was to increase the concentration of retinoic acid (RA) by 5-fold (5µM). A recent study addressed variability in the populations of terminally differentiated neurons from multiple standard protocols.^24^ It showed that including growth factors bFGF and FGF-8, along with Noggin during the neural induction increased the viability of neural progenitors, and resulted in an optimized expression of the desired neural populations. An unexpected advantage of this approach was the formation of smaller EBs, which we speculate helped with the diffusion of RA, the transcription factor responsible for neuronal differentiation. The efficacy of this protocol revision was validated using RT-qPCR and ultimately resulted in the upregulation of genes related to glutamatergic neurotransmission^25^, GRIA1 (1.41-fold; p = 0.011), GRIA2 (1.47-fold; p = 0.017) and GRIN2A (1.35-fold; p = 0.031), each respectively correlated to the expression of AMPA receptor subunit 1 and subunit 2, and the NMDA receptor subunit 2A (**Fig 2b**). Moreover, we observed a significantly higher upregulation for SLC17A7 (2.31-fold, p<0.001), a gene correlated with the expression of glutamate vesicles. On the other hand, we did not observe any significant change in the expression of genes related to synaptic transmission, NLGN (0.98-fold, p = 0.51) and NLG4 (1.08-fold, p = 0.244)^26^ nor statistically significant changes in the expression of genes related to neuronal maturation, RBFOX3 (0.98-fold, p = 0.531) and MAP2 (1.01-fold, p = 0.71). These observations point to an up regulation of glutamatergic neurons with respect to the total of the neuronal population or the number of synapses. Unexpectedly, we observed a significant upregulation in the glial markers, CNPase (1.75-fold, p < 0.001) and GFAP (2.33-fold, p < 0.001).

**Figure 2.**
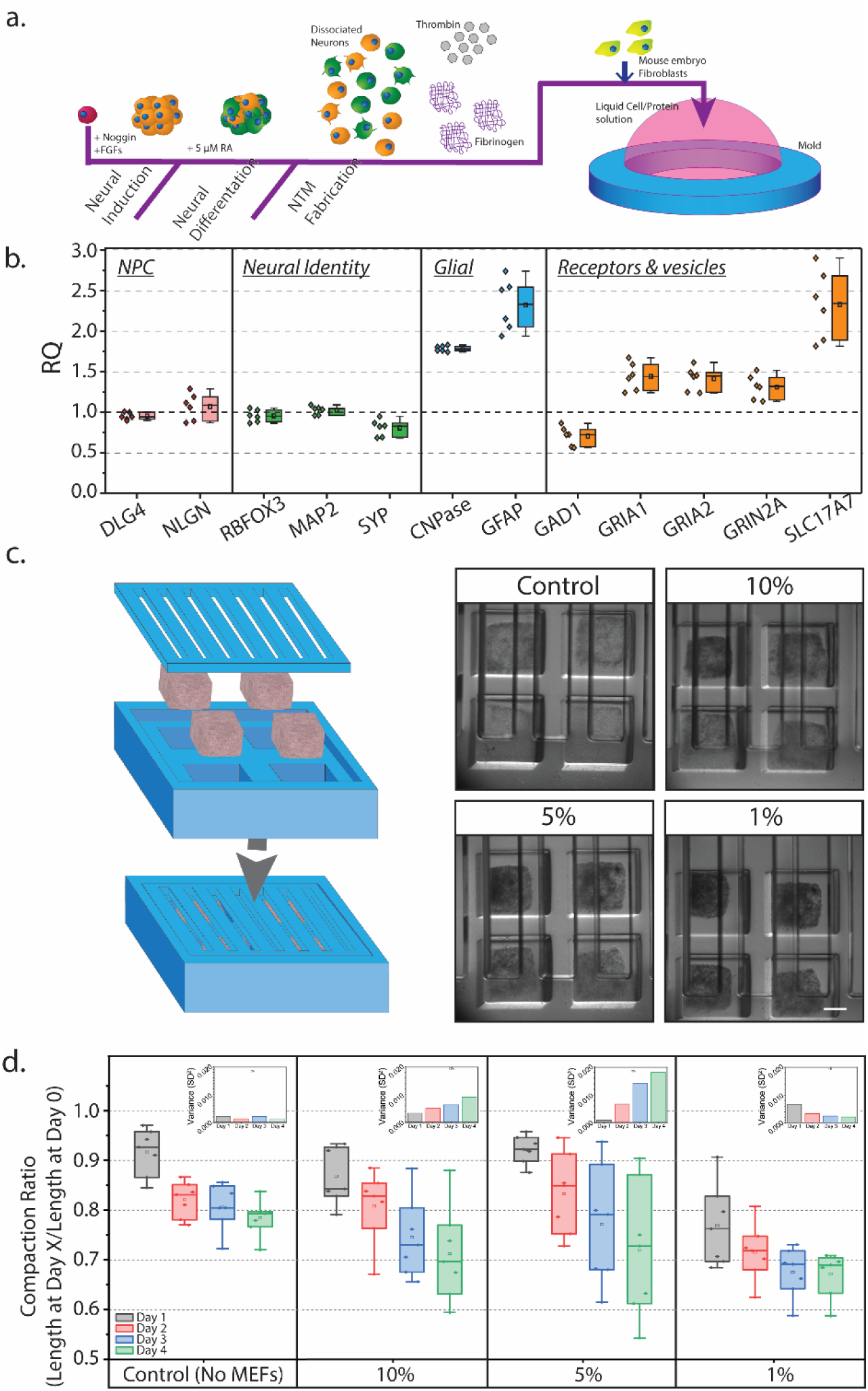
Adding MEFs facilitates the compaction of NMT enriched with glutamatergic neurons. **a.** Diagram representing the on-chip fabrication protocol. **b. c.** Box plots showing RQ values for RT-qPCR done on key genes of interest normalized for standard differentiation protocol versus the tested enriching differentiation protocol color coded in relation to functionality (pink: neural development; green: neuronal identity, blue: glial cells in interest, orange: neurotransmitters and receptors of interest) (n=6, *p<0.05 T-test) **c.** Representative images of resulting compaction after 4 DIV at different ratios of MEF to enriched neural progenitors. (Scale bar: 500 µm) **d.** Box plots showing the compaction ratio (compared between day and day 0) for 4 DIV at the different ratios of MEF to enriched neural progenitors. (n = 4, *p<0.05 ANOVA with post-hoc Tukey test)

Unfortunately, alteration of the standard differentiation protocol used in prior work^27^ resulted in cell-protein constructs that would show high cell viability but failed to compact and achieve the necessary tensile strength to sustain fluid shear stress. This was addressed by adding mitotically arrested mouse embryonic fibroblasts (MEFs) to the liquid cell-protein solution prior to aliquoting to their respective molds. Fibroblasts have high traction forces^28^, and their mitotic inactivation would allow us to add an aid for compaction without introducing a cell type that would overtake the neuronal population. We tested different ratios of fibroblasts to dissociated neurons on their effect on the compaction of the neural tissue mimics (**Fig 2c**). We observed that adding 10% or 5% of the total number of neuronal cells as MEFs showed a trend of improvement (ANOVA with Tukey: p = 6.3E-2, 95% C.I. = [−9.9E-2,1.8E-2]; p = 7.1E-1, 95% C.I. = [−7.1E-2,3E-2] respectively) in compaction ratio – length at day 4 after fabrication normalized by length at day of fabrication (day 0) – when compared with the compaction ratio of the control samples (**Fig 2d**). On the other hand, the 1% samples showed a statistically significant decrease in the compaction ratio when compared to control samples (ANOVA with Tukey: p = 1.25E-9, 95% C.I. = [−1.7E-1, −7.4E-2]). Moreover, we also observed that the 10% and 5% samples’ sizes increased in variability during the first four days of compaction, while the samples with 1% MEFs resulted in constructs with variability notably smaller than the 10% (3.75-fold) and 5% (9.03-fold) samples. While unexpected, these observations might indicate a technical limitation in mixing the MEFs homogenously when adding them during the fabrication phase (day 0) without harming the dissociated neurons. Therefore, we decided on adding MEFs at 1% the total amount of EB-derived neuronal cells during the biofabrication of the NTMs.

### On-chip fabrication of NTM requires an initial neuron monolayer to achieve good connectivity

One of the benefits reported for hydrogel-neuronal fabrication of neural tissue mimics is their compatibility with different testing platforms while still achieving wide-ranging connectivity throughout the structure^27^. However, during initial testing of the molding tissues directly on the MEA (T), we failed to record any network spike trains. We hypothesized that, during compaction, neurons suspended in the hydrogel would migrate to other neurons. For many neurons this would mean displacing away from the region of detection at the surface of the electrodes. To mitigate this issue, we hypothesized that adding a monolayer (ML) of dissociated neurons would ensure electrical connection, while simultaneously attracting the neurons engrafted in the hydrogel (**Figure 3a**). Confocal imaging showed that the cell distribution within the tissue was very heterogenous (**Fig 3b.i**) in addition to a visible (~50µm) gap between the bottom of the substrate and the first cluster of neurons embedded in the hydrogel (**Fig 3b.ii** and **Fig 3b.iii**). When we performed imaging on a neural monolayer, we confirmed that our high-density seeding protocol maintained a homogenous distribution of cells (**Fig 3c.i**) for at least 3 DIV, and, as expected, good adhesion to the substrate (i.e. good contact with electrodes; **Fig 3c.ii & iii**). It was hypothesized that a higher than seeding density, as usually reported in literature^29^, was important to ensure a homogenous adhesion layer for the tissue fabrication stage. Therefore, twelve to twenty-four hours prior to tissue fabrication, a sample of EBs would be dissociated and seeded on the MEA substrate at a density of 1E5 cells/mm^2^. It would be then incubated in NDM supplemented with 0.1 mg/mL of Matrigel™ until it was time to seed the cell-protein solution. After fabricating the tissue on top of the monolayer (ML+T), confocal imaging showed a more uniform cell distribution within the construct (**Fig 3d.i**). It was also notable that there was no gap between the substrate and the first layer of cells and there were neurites extending in the z-direction throughout the ~60 µm that confocal microscopy was able to image (**Fig 3d.ii** and **3d.iii**).

**Figure 3.**
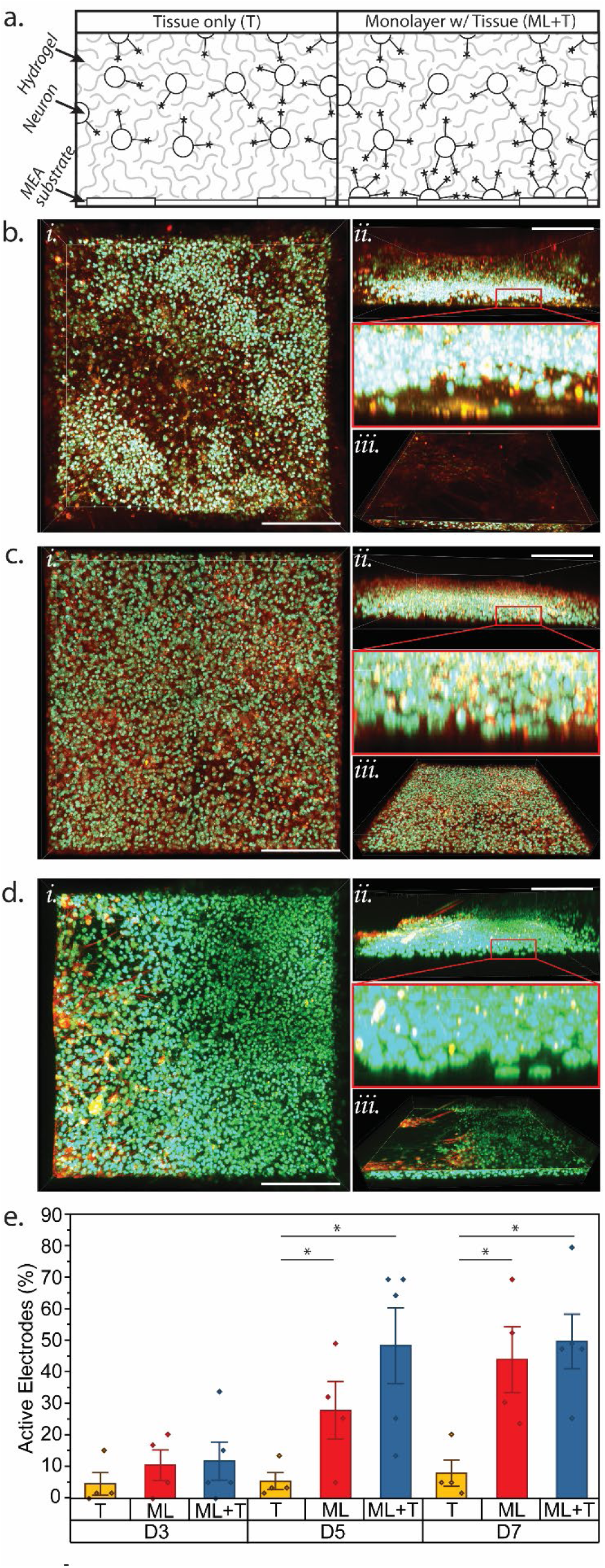
Fabricating NTMs on neuronal monolayer increases recording efficiency. **a.** Representative diagram showing how the absence of a monolayer prior to neural tissue fabrication results in a tissue compaction that fails to make contact with electrodes. Confocal images were taken to show the cell distribution (*i*), the side profile (*ii* & *iii*) and the bird’s eye view (*iv*) of tissue only (**b**), monolayer only (**c**) and tissue on top of monolayer (**d**) fabricated directly on the MEA. (scale bar: 100µm) **e.** Bar plot showing the percent of active electrodes between MEA with NTM only (T), monolayer only (ML) and NTM fabricated over a monolayer (ML+T) on days 3, 5 and 7 *in vitro.* (n = 5, *p<0.05 ANOVA with post-hoc Tukey test)

Recordings showed that, by day 5 of culture on the MEAs, the percentage of active electrodes, i.e. electrodes that were detecting at least 10 spikes/min, was larger for ML samples (ANOVA with Tukey: p = 1.5E-2; 95% C.I. = [−39.36, −3.58]) and for ML+T samples (ANOVA with Tukey: p = 2E-4; 95% C.I. = [−47.62, −13.68]) than for control T samples (**Fig. 3e**). There was a trend of ML+T samples showing a higher increase in active electrodes by D5, which we hypothesized to coincide with a larger number of synapses being formed with the addition of the NTM. This hypothesis was supported by general action potential firing parameters (**Supp. Figure 4**). For ML+T samples, we observed a transient higher increase (compared to ML samples) in firing rate prior to settling into a steady state (**Supp. Figure 5**), as it is common to see during the later phases of synaptogenesis of *in-vitro* neuronal cultures.

### Patterned stimulation effectively modulates firing patterns in ML+T samples

Given the thickness of the NTM, we tested the effectiveness of patterned stimulation from the Polygon1000 system to evoke responses by depolarizing the neurons throughout the thickness, (as opposed to indirectly evoking a response by only stimulating the neurons at the NTM boundary that innervate neurons in contact with electrodes). To do this, we stimulated NTMs with flood exposure at 250/500/750/1000 W using a 1s stimulation trains at 50Hz (**Fig. 4a**). Recordings showed that stimulations evoked properly timed responses at 250W (**Fig. 4b**). Regardless, there was an increase in evoked responses that seems to have saturated at 750W, showing a slight average decrease in response from the samples when stimulated at 1000W. Given that optogenetic activation is not hindered by using excess intensity, this decrease in response could be a limitation in the system’s response for the 50Hz frequency, not allowing the ChR2 channels to fully close. Moreover, because high output power can cause tissue damage, we designed our experiments using the 750W intensity with the system.

**Figure 4.**
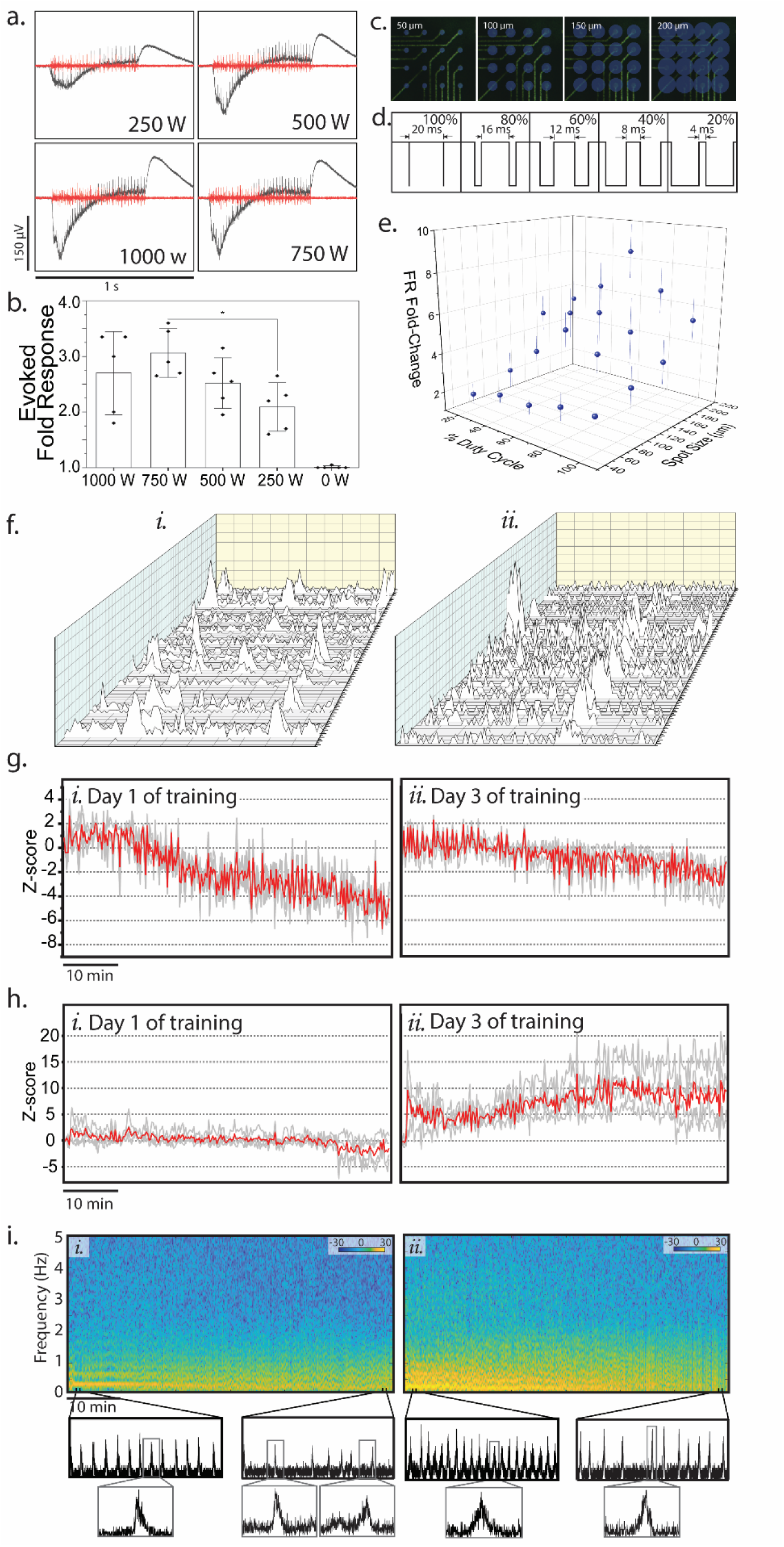
Incorporating tissue-wide spatial optogenetic stimulation can modulate large scale firing patterns of engineered NTMs. **a.** Representative exports of raw (black) and high-pass filtered (red) of recordings during stimulation trains at different intensities (*i.* 0 W, *ii.* 250 W, *iii.* 500 W, *iv.* 750 W and *v.* 1000 W). **b.** A bar graph shows the evoked fold response to stimulation at different intensities with respect to firing rate prior to stimulation. (n = 6, *p<0.05 ANOVA with post-hoc Tukey test). **c.** Representative image of electrodes with the respective spot sizes (*i.* 50 µm, *ii.* 100 µm, *iii.* 150 µm and *iv.* 200 µm) compared along with for duty cycles that were compared (**d.**). **e.** 3D graph showing how spot size and % duty cycle affected firing rate fold-change (n=6). **f.** Representative cascade plots of binned spikes for MEA electrodes between day 1 (*i.*) and day 3 (*ii.*) **g.** Z-score of spike firing rate throughout the 60 minutes of recording for day 1 (*i*) and day 3 (*ii*). Shaded gray line plots show individual sample recordings, and red line plot represents the average at each time point. **h.** Z-score of burst firing rate throughout the 60 minutes of recording for day 1 (*i*) and day 3 (*ii*). Shaded gray line plots show individual sample recordings, and red line plot represents the average at each time point. **i.** Spectrogram for recording done on training day 1 (*i*) and training day 3 (*ii*) of training. Excerpts of corresponding binned spikes summed for all recording electrodes for that time point pertaining to minute 5 and minute 55 of recording. Below zoom-ins of single spike trains.

Furthermore, we tested the NTM’s response to different stimulation duty cycles as well as spot sizes. Spot sizes of 50µm (corresponding to a single electrode diameter), 100µm, 150µm and 200µm (corresponding to approximately flood exposure) (**Fig. 4c**) were tested against duty cycles ranging from 20-100%. **Figure 4d** shows the corresponding pulse widths for the tested duty cycles. We desired a duty cycle that would elicit the highest sensitivity to different stimulation patterns. As expected, the smallest spot size didn’t noticeably change with duty cycles as thin beams would likely be easier to refract as it travelled through the tissue. However, across the remaining spot sizes, there was a significant increase in firing rate fold change between the 60% duty cycle (corresponding to a pulse width 12ms) and the rest. This consistent trend indicated the duty cycle at which we could achieve the highest evoked activation prior to reaching the range that would fatigue the ChR2, which results in decreased optogenetic efficiency. These observations also informed our design of stimulation patterns having at least a width thickness of 100µm, as it would allow for the smallest size that would be able to elicit a noticeable response for the 3D NTMs.

After determining the performance constraints of our system, NTM samples on chip were fabricated and allowed 4 days on chip to compact prior to being subjected to a training regimen. A diagonal beam would be shone on the sample starting at the bottom right of the sample and then progressed in a step-wise fashion until the top-left corner of the sample is reached (**Supp Figure 6**). This pattern would be repeated daily for an hour while recordings were done simultaneously. An enclosure around the MEA samples along with HEPES buffer was used to control pH change during this time without disrupting the optogenetic stimulation^30^. The effectiveness of HEPES buffer was monitored by observing the hue of the phenol red in the culture media, and ensuring there was no observable change in media color.

While these tissues have shown connectivity throughout the construct in previous studies^27^, a desired and expected observation from the training regimen was to observe a higher synchronicity in bursting between the trained samples (**Fig. *4fii***) than the untrained samples (**Fig. 4f*i***) Furthermore, we quantified the changes in firing throughout the training process with respect to the baseline by using the z-score of the spiking rates (**Fig. 4g**). Baseline measurements were obtained from the first 5 minutes of stimulation (not shown) to discount effects from the general response to optogenetic stimulation, thus focusing on the effects of the prolonged exposure of the directional stimulation pattern. In the first day of training (**Fig. 4g*i***), NTM samples continued to show a transient increase in firing rate for approximately the first 15 minutes of stimulation, proceeded by a steep decrease in z-score for the remainder of the stimulation process. Moreover, the removal of the stimulation patterns seemed to have caused an additional decrease in spiking rate during the post-stimulation spontaneous recordings.

These observations coincide with our hypothesis of weakening established synapses by using the directionality of the transient stimulation patterns. On day three of training (**Fig. 4g*ii***), spiking rate did not show this continued increase in response, but rather stayed stable through the first half of the recordings, before showing a slight drop off on the second half. This observation might be more indicative of neuronal fatigue exposed to prolonged stimulation. Long term harm on the cultures were dismissed by the observation of recovery of normal firing rates upon the following day of recording.

Analogous measurements were done on the bursting rate in the recordings (**Fig. 4h**). For these measurements, bursting events were defined based on network bursting events. Bursting events are usually defined by a fixed set of constraints related to a predetermined number of spikes per burst, minimum burst duration, and a limit in the spacing between individual spikes within a prospective burst, among others. Given the dynamic nature of the study, this definition lacked the specificity needed to observe nuanced progression of the bursting events. For our purposes, bursts were based on network wide (defined as at least affecting 1/6^th^ of the active electrodes) spike trains. Spike train widths were defined using the sum of the raster plots across all channels and binned in 10ms time intervals. The widths of resulting firing peaks were then averaged. Bursts in individual electrode recordings were defined as spike trains that were at least one standard deviation below the average width of the network spikes described above.

The z-score for these bursting events remained statistically unchanged for the majority of the recording session, with two samples showing a transient increase at the beginning of the recording, as well as a decrease when the stimulation regimen was removed. Interestingly, at day 3 of training, the bursting z-score showed a noticeable upward drift, with no noticeable change after the stimulation was removed. These observations were interpreted as the neural network responding to the training regimen differently based on whether the regimen was a familiar stimulus to the network.

On day 1 of training, a steep decrease in spiking z-score with a steadily unchanging bursting z-score could be indicative of a decrease in burst size along with a general decrease in single spike firing rate while the bursting rate remains unchanged. This was a stark contrast with the observations for day 3 of training, where while there was a smaller decrease in spiking rate from baseline, on average, the bursting z-score increased. This in turn would be indicative of restructured firing behavior, with single spikes decreasing and firing mainly occurring within bursts. This was validated by qualitative observation of the raster plots (**Supp. Figure 7**).

Power spectra density (PSD) plots were obtained from the binned summations across channels per day of recording described above. It was noticeable that prior to perturbation, untrained networks showed clear harmonics in the spectra, which became disrupted throughout the training regimen (**Fig. 4i.i**). Harmonics in a PSD of a neurological signal is a sign of periodicity given the non-sinusoidal nature of the waveforms that result from the summation of the electrode recordings. Interestingly, for the last day of training this periodicity was recovered but with a loss in the smoothness of the PSD signal, which would indicate a network firing at an approximate periodicity with a measurable variability (**Fig. 4i.ii**). This periodicity was also completely disrupted for later times in the recording as the training progressed.

These changes in periodicity can also be qualitatively appreciated in excerpts of the signal from the corresponding time intervals from the PSD plots. However, looking closer at the individual peaks, their shapes strongly validate how the stimulation regimen had a sustained effect on the timing of the bursting events recorded in each individual channel. Prior to any training, the beginning of these peaks shows a synchronous start of the bursting events followed by a gradual decline. While some peaks at the end of the first training day maintained this shape, other peaks during the same time interval showed a more gradual rise, followed by a steeper fall. Moreover, at the beginning of the last day of training, peaks exhibited a similar gradual rise followed by a gradual drop, later turning into a steep drop consistently at the end of the training regimen.

### Patterned stimulation imparts directionality in the networks bursting

We quantified the effect of our training regimen on evoking directionality in the firing by first ordering the recording channels in the raster plots with respect to the direction of the stimulation patterns. Directionality would be observed as a shift in the timing of bursting events across the different ordered channels. When ordering the electrodes for recording from untrained networks, a magnified observation of a network-wide bursting events shows synchronicity but no discerned directionality (**Fig. 5a**). To quantify this observation, individual channel recordings were binned by time intervals equal to half of the calculated burst width as explained for **Figure 4h**.

**Figure 5.**
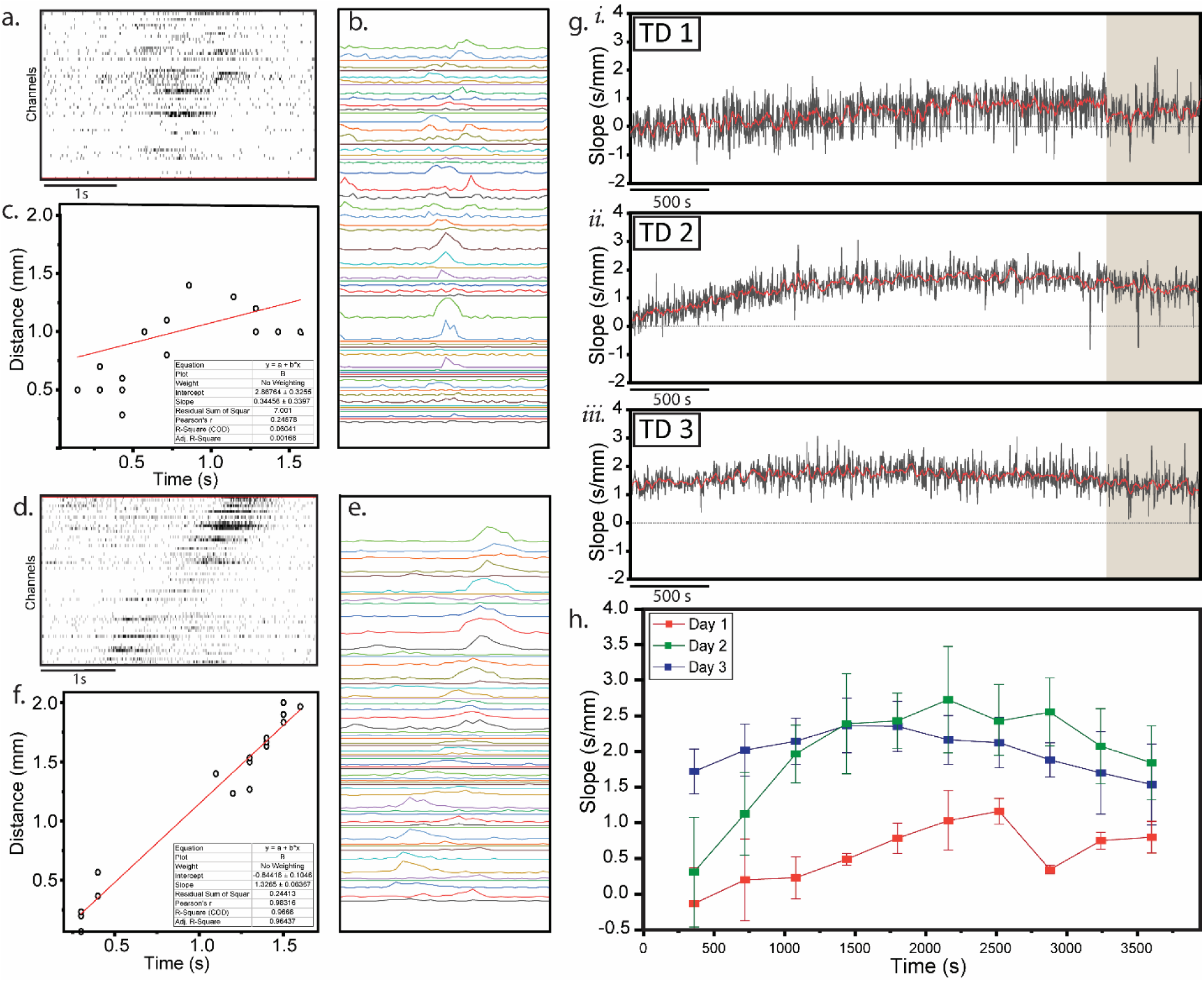
Tissue-wide spatial stimulation can evoke directional firing patterns in large-scale engineered NTMs. **a.** Raster plot of a network wide bursting event at 60s (start of training) and 3000s (end of training) of recording for training day 1 (TD 1) **b.** Binned spikes representing peaks corresponding to bursting events. **c.** Linear fit for the time points of the local maxima from **b** plotted against the position corresponding to the respective electrodes. **d, e** and **f** replicate **a, b** and **c** for training day 3 (TD 3). **f.** Line plot showing the resulting slope of each network wide bursting event throughout the recording for training day 1, 2 and 3 (TD 1 (*i*), TD 2 (*ii*) and TD 3 (*iii*)) for a representative sample. Brown shading corresponds to the time the optogenetic stimulation was removed. **c.** Line plot of the mean slope for 600s bins and averaged across 3 experimental samples for day 1, day 2 and day 3 of training.

Electrodes that burst also showed peaks whose local maxima was extracted for each detected network wide burst (**Fig. 5b**). The time location of the local maxima was plotted with respect of the electrode location. A linear fit for these scatter plots resulting from untrained networks showed slope calculations that were not statistically different from zero (**Fig. 5c**).

In contrast, after training, close-up visualizations of network bursting events showed a progressive time shift in the respective bursts for each channel in the direction of the stimulation (**Fig. 5d**). Binning the spike trains as described above showed peaks that further illustrate the gradual shift in the local maxima (**Fig. 5e**). The linear fit for the scatter plot for the time shift of the peaks against the distance across the electrode grid showed an R^2^ value of ~0.97 and a Pearson’s r value of 0.98 (**Fig. 5f**). To see the real-time progression of the slope values as a result of the training, the slope of each network bursting event was calculated and plotted against time of recording (**Fig. 5g**). The resulting line plot for training day 1 (TD 1), as expected, started with slopes oscillating around zero prior to showing a gradual shift in the positive direction as the training progressed. When the stimulation was removed, a sudden drop can be observed. At TD 2, the slope profile again seemed to start around zero, but increased at a faster rate than in TD 1. Surprisingly, there seemed to be very little change during the post-training spontaneous phase of the recording. This positive slope phase seemed to be sustained all throughout TD 3, staying sustained during both recording of the training and post-training phases. The respective R^2^ values for each corresponding fit can be observed in **Figure 5c, e**.

These observations were quantified and summarized by binning the slope values for every 6 minutes of recording and averaged across samples (**Fig. 5h**). The means followed a similar trend as presented in **Figure 5g**. TD 1 showed samples having near-zero slopes which responded to stimulation by progressively increasing prior to having a steep drop in response to the removal of the training regimen. Slope values on TD 2 consistently showed a steep increase for the first 20 minutes of training and were sustained for the remainder of the training, followed by a gradual decrease when the training was removed as opposed to the sudden decrease observed for TD 1. Furthermore, TD 3 consistently had positive slopes yet started a decrease prior to the removal of stimulation and presented no detectable response to the removal of stimulation. This might be due to an adverse response to a stimulation trying to reinforce synapses that have reached a limit in potentiation, thus additional evoked bursts cause no further potentiation^31^.

## Conclusion

As more methods arise for the engineering of tissues that mimic neural behavior, efforts should be invested towards also being able to modulate the firing patterns of the resulting constructs. Expanding both the capability of engineering the shape of these NTMs as well as training their behavior will continue to lay the foundation for the development of more sophisticated neuromorphic networks^32^. In this work, we showed the ability to establish directionality across a mm-scale on-chip fabricated 3D NTMs by timed stimulation that mimicked STDP driven plasticity. Close-loop stimulation has been showing effectiveness to train networks at the synaptic scale^33^. By the enrichment of key neural populations, and a compatible training regimen, we show that this method can also be effective to modify firing patterns at a larger scale. This platform lends itself for further exploration of the mechanistic pathways that lead to this modulation as well as expanding its implementation to more complex functional outputs that can take advantage of a diversity of NTM morphologies.

## Methods

### mESC culture and differentiation

Mouse-derived embryonic stem cells (mESC) were cultured and differentiated as has been presented in previous publications^19, 27^. Briefly, optogenetic (ChR2::TdTomato) mouse embryonic stem cells (mESC) were expanded on top of a feeder layer consisting of mitomycin-C inactivated mouse embryonic fibroblast in standard mESC expansion media. Upon confluency of colonies, stem cells were dissociated from fibroblasts and resuspended in low adhesion petri dishes in Neural Induction Media consisting of 1:1 cocktail solution of Advanced DMEM/F12 (Life Tech) and Neurobasal (Life Tech), 10% KnockOut Serum (Invitrogen), 1% Pen-Strep (Invitrogen), 1% GlutaMax (Invitrogen), 0.1 mM of β-mercaptoethanol (EMD Millipore) supplemented with 1% Non-essential amino acids (Invitrogen), 50ng/mL of Noggin (Invitrogen), 20 ng/mL of bFGF (Invitrogen) and 20 ng/mL FGF-8 (Invitrogen). After two days, formed embryoid bodies were resuspended in media now supplemented with 2.5% B-27, 1% N2-Supplement (Invitrogen), 1% ITS Supplement-B (EMD Millipore), 100 µM Ascorbic Acid (EMD Millipore), 0.2% Heparin (Sigma-Aldrich), and 5µM Retinoic Acid (Sigma-Aldrich) (Neural Induction Media).

### Tissue formation

On day 5 of differentiation, neurospheres were dissociated in Accutase (EMD Millipore) and seeded on functionalized custom-made MEAs or glass substrates at ~5000 cells/mm^2^ and incubated (37°C/5% CO_2_) in Neural Differentiation Media supplemented with 1 µg/mL of Matrigel™ for 12-24 hrs. This showed to improve adhesion with the glass surface and the subsequent incorporation of the 3D neural tissue. The tissue was formed directly on top of the neuronal monolayer following previous protocol^27^, using a 12.5e6 cell/mL solution of dissociated neural spheres with 30% Matrigel (Corning), 4 mg/mL, fibrinogen (Sigma-Aldrich), and 2 U/mg_Fibrinogen_ of thrombin (Sigma-Aldrich) that spontaneously cross-linked to form the 3D neural tissue mimic. Control tissue mimics were seeded on functionalized MEAs that had been incubated with only 1ug/mL of Matrigel. These constructs were subsequently cultured in Neural Differentiation Media supplemented with 10 ng/mL each of BDNF, GDNF and CNTF (Neuromics) for 4 days during compaction. For the remainder of the experiments, constructs were maintained in Neural Maintenance Media consisting of Neurobasal supplemented with 2.5% B-27(Gibco), 1% Pen-Strep (Invitrogen), 1% GlutaMax (Invitrogen), 0.1 mM of β-mercaptoethanol (EMD Millipore) and 10 ng/mL of BDNF.

### Electrophysiology recording and optogenetic stimulation

NTMs and monolayer samples assigned for recording were cultured in in-house fabricated customized Pt-black microelectrode arrays made on borofloat glass substrates following standard lithographic techniques^19^. Electrophysiological recordings were done using an MEA 2100 Lite Amplifier (MultiChannel Systems MCS GmbH Germany) with the heating plate kept at 37°C. Measurements were performed in dark at a sampling rate of 10kHz. Spontaneous recordings were done within CO_2_ chambers. For cultures undergoing recordings during optogenetic stimulation, media was supplemented with 25mM HEPES buffer to control for pH changes. This concentration was selected empirically by assessing the lack of color change in Phenol Red after 45mins in a controlled environment. CO_2_ chambers were not applicable for recordings undergoing stimulation due to refraction caused by the glass chamber lid.

Stimulation was performed with the Polygon1000-G/Large-field of view (Mightex, Pleasanton CA) at 465nm. This system was fitted into a customized Nikon Eclipse FN1 (Nikon, Melville NY) using a 10x objective to achieve single-digit micrometer pixel size of incident light at high fps within physiologically relevant temporal ranges. Pattern sequences of stimulation were produced in MATLAB.

### Immunostaining and imaging

Immunostaining was performed on samples cultured on glass substrates after being fixed with 4% paraformaldehyde and treated with 0.05% Triton-X. Permeabilized samples with blocked with 2% bovine serum albumin in PBST at 4°C overnight. Samples were later stained with primary antibodies at 4°C overnight, followed by staining of secondary antibodies at room temperature for 2 hrs. All antibodies were diluted in PBST. The used antibodies were the following: anti-tubulin III polyclonal mouse antibody (Abcam), anti-MAP2 polyclonal rabbit antibody (Abcam), and anti-synaptophysin 1 polyclonal chicken antibody (Thermofisher). Samples were then counterstained, respectively, with goat anti-tubIII IgY AlexaFluor 488 (Thermofisher), goat anti-MAP-2 IgY AlexaFluor 568 (Thermofisher), goat anti-chicken IgY AlexaFluor 647 (Thermofisher), respectively, along with DAPI as a nuclear stain. After washing overnight at 4°C in blocking buffer, samples were mounted with Prolong Diamond antifade (ThermoFisher, MA) and imaged using Zeiss 710 Confocal microscope (Carl Zeiss Microscopy).

### RT-qPCR

Biofabricated NTMs designated for RT-qPCR were allowed to compact and mature for a week in suspension prior to being individually flash frozen and stored in −80°C until mRNA extraction. Total RNA was collected and purified using the RNeasy Mini Kit Part 1 (Qiagen). The total RNA concentration was quantified using a Nanodrop spectrophotometer. Only samples with absorption ratios (A260/280) ~2.0 were included in this study. Synthesis of cDNA was done with 100ng RNA per sample with SuperScript IV VILO EzDNase Kit (ThermoFisher Scientific). Triplicate assays were performed by QuantStudio 3 (ThermoFisher Scientific) in TaqMan Universal Master Mix II (Applied Biosystem). The probes used for monitoring development and enrichement of neuronal populations for this study were: NMDAR1 (Glutamanergic Neurons) [Assay ID: Mm00433790_m1], GAD67 (GABAergic Neurons) [Assay ID: Mm04207432_g1], GFAP (Astrocytes) [Assay ID: Mm01253033_m1], CNPase (Myelinating Oligodendrocytes) [Assay ID: Mm01306641_m1]. Glyceraldehyde3-phosphate dehydrogenase (GAPDH, assay ID: Mm99999915) was used as the endogenous reference gene. RT-PCR was run at 50 °C for 2min, 95 °C for 2min, for 40 cycles [Denaturing: 95 °C for 1 s, Extend/Annealing: 60 °C for 20 sec] with ROX, FAM and MGB as reference dye, reporter, and quencher, respectively.

## Acknowledgements

This study was funded by NSF EFRI C3 SoRo No. 1830881, NSF Expeditions: Mind in Vitro — Computing with Living Neurons, No. 2123781, and the Strategic Research Initiatives (SRI) Program of the University of Illinois at Urbana-Champaign, as well as the National Science Foundation Science and Technology Center Emergent Behavior of Integrated Cellular Systems Grant No. 3939511 and National Science Foundation Research Traineeship-Miniature Brain Machinery Grant No. 1735252. We thank Dr. Roger Kam from Massachusetts Institute of Technology who provided the transfected cell line used in this study. We are grateful to the staff at the Carl R. Woese Institute for Genomic Biology for their support as well as Dr. Glennys Mensing and Joseph Maduzia for assisting in the fabrication of the microelectrode arrays.

## Author contributions

G.J.P. and R.B. designed the experiments, G.J.P performed data acquisition and analysis of experiments, and writing of the manuscript; M.W., E.K., S.R., O.A., Y.K., L.G. and M.S.A.T. contributed with data acquisition; D.B. contributed with PCR data acquisition and analysis; R.B. led the project and edited the manuscript.

## References

1. Prochazka, A. Neurophysiology and neural engineering: a review. J. Neurophysiol. 118, 1292–1309 (2017).

2. Aydin, O. et al. Neuromuscular actuation of biohybrid motile bots. Proc. Natl. Acad. Sci. 116, 19841–19847 (2019).

3. Aydin, O. et al. Development of 3D neuromuscular bioactuators. APL Bioeng. 4, 016107 (2020).

4. Ko, E. et al. Matrix Topography Regulates Synaptic Transmission at the Neuromuscular Junction. Adv. Sci. 6, 1801521 (2019).

5. Osaki, T., Uzel, S. G. M. & Kamm, R. D. On-chip 3D neuromuscular model for drug screening and precision medicine in neuromuscular disease. Nat. Protoc. 15, 421–449 (2020).

6. Wagenaar, D. A., Pine, J. & Potter, S. M. Searching for plasticity in dissociated cortical cultures on multi-electrode arrays. J. Negat. Results Biomed. 5, 16 (2006).

7. Wheeler, B. C. & Brewer, G. J. Designing neural networks in culture. Proc. IEEE 98, 398–406 (2010).

8. Ide, A. N., Andruska, A., Boehler, M., Wheeler, B. C. & Brewer, G. J. Chronic network stimulation enhances evoked action potentials. J. Neural Eng. 7, 16008 (2010).

9. Joo, S., Song, S. Y., Nam, Y. S. & Nam, Y. Stimuli-responsive neuronal networking via removable alginate masks. Adv. Biosyst. 2, 1800030 (2018).

10. Massobrio, P., Tessadori, J., Chiappalone, M. & Ghirardi, M. In-vitro studies of neuronal networks and synaptic plasticity in invertebrates and in mammals using multielectrode arrays. Neural Plasticity https://www.hindawi.com/journals/np/2015/196195/ (2015) doi:10.1155/2015/196195.

11. George, R. et al. Plasticity and Adaptation in Neuromorphic Biohybrid Systems. iScience 23, 101589 (2020).

12. Tessadori, J., Bisio, M., Martinoia, S. & Chiappalone, M. Modular neuronal assemblies embodied in a closed-loop environment: toward future integration of brains and machines. Front. Neural Circuits 6, (2012).

13. DeMarse, T. B., Wagenaar, D. A., Blau, A. W. & Potter, S. M. The neurally controlled animat: biological brains acting with simulated bodies. Auton. Robots 11, 305–310 (2001).

14. Novellino, A. et al. Connecting neurons to a mobile robot: an in-vitro bidirectional neural interface. Comput. Intell. Neurosci. 2007, 1–13 (2007).

15. Feldman, D. E. The Spike-timing dependence of plasticity. Neuron 75, 556–571 (2012).

16. Buonomano, D. V. & Carvalho, T. P. Spike-Timing-Dependent Plasticity (STDP). in Encyclopedia of Neuroscience (ed. Squire, L. R.) 265–268 (Academic Press, 2009). doi:10.1016/B978-008045046-9.00822-6.

17. Waters, J., Nevian, T., Sakmann, B. & Helmchen, F. 4.39 - Action Potentials in Dendrites and Spike-Timing-Dependent Plasticity. in Learning and Memory: A Comprehensive Reference (ed. Byrne, J. H.) 803–828 (Academic Press, 2008). doi:10.1016/B978-012370509-9.00029-2.

18. Shouval, H., Wang, S. & Wittenberg, G. Spike Timing Dependent Plasticity: A Consequence of More Fundamental Learning Rules. Front. Comput. Neurosci. 4, (2010).

19. Pagan-Diaz, G. J. et al. Modulating electrophysiology of motor neural networks via optogenetic stimulation during neurogenesis and synaptogenesis. Sci. Rep. 10, 12460 (2020).

20. Chuang, J.-H., Tung, L.-C., Lee-Chen, G.-J., Yin, Y. & Lin, Y. An approach for differentiating uniform glutamatergic neurons from mouse embryonic stem cells. Anal. Biochem. 410, 149–151 (2011).

21. Chuang, J.-H., Tung, L.-C., Yin, Y. & Lin, Y. Differentiation of glutamatergic neurons from mouse embryonic stem cells requires raptor S6K signaling. Stem Cell Res. 11, 1117–1128 (2013).

22. Lin, Y. Differentiation of Embryonic Stem Cells into Glutamatergic Neurons (Methods). in Stem Cells and Cancer Stem Cells, Volume 6: Therapeutic Applications in Disease and Injury (ed. Hayat, M. A.) 47–55 (Springer Netherlands, 2012). doi:10.1007/978-94-007-2993-3_5.

23. Wichterle, H. & Peljto, M. Differentiation of mouse embryonic stem cells to spinal motor neurons. in Current Protocols in Stem Cell Biology (eds. Bhatia, M. et al.) (John Wiley & Sons, Inc., 2008). doi:10.1002/9780470151808.sc01h01s5.

24. Wu, C.-Y., Whye, D., Mason, R. W. & Wang, W. Efficient Differentiation of Mouse Embryonic Stem Cells into Motor Neurons. J. Vis. Exp. (2012) doi:10.3791/3813.

25. Meadows, J. P. et al. DNA methylation regulates neuronal glutamatergic synaptic scaling. Sci. Signal. 8, ra61 (2015).

26. Ricceri, L., Michetti, C. & Scattoni, M. L. Chapter 17 - Mouse Behavior and Models for Autism Spectrum Disorders. in Neuronal and Synaptic Dysfunction in Autism Spectrum Disorder and Intellectual Disability (eds. Sala, C. & Verpelli, C.) 269–293 (Academic Press, 2016). doi:10.1016/B978-0-12-800109-7.00017-0.

27. Pagan-Diaz, G. J. et al. Engineering geometrical 3-dimensional untethered in vitro neural tissue mimic. Proc. Natl. Acad. Sci. 116, 25932–25940 (2019).

28. De La Pena, A., Mukhtar, M., Yokosawa, R., Carrasquilla, S. & Simmons, C. S. Quantifying cellular forces: Practical considerations of traction force microscopy for dermal fibroblasts. Exp. Dermatol. 30, 74–83 (2021).

29. Wongpaiboonwattana, W. & Stavridis, M. P. Neural Differentiation of Mouse Embryonic Stem Cells in Serum-free Monolayer Culture. J. Vis. Exp. JoVE 52823 (2015) doi:10.3791/52823.

30. Will, M. A., Clark, N. A. & Swain, J. E. Biological pH buffers in IVF: help or hindrance to success. J. Assist. Reprod. Genet. 28, 711–724 (2011).

31. Kilborn, K., Lynch, G. & Granger, R. Effects of LTP on response selectivity of simulated cortical neurons. J. Cogn. Neurosci. 8, 328–343 (1996).

32. Kagan, B. J. et al. In vitro neurons learn and exhibit sentience when embodied in a simulated game-world. 2021.12.02.471005 Preprint at https://doi.org/10.1101/2021.12.02.471005 (2021).

33. Chen, R., Canales, A. & Anikeeva, P. Neural recording and modulation technologies. Nat. Rev. Mater. 2, 1–16 (2017).

