## Supplementary Figures for "Engineering 3D neuronal networks with directional endogenous neuronal plasticity pathways"

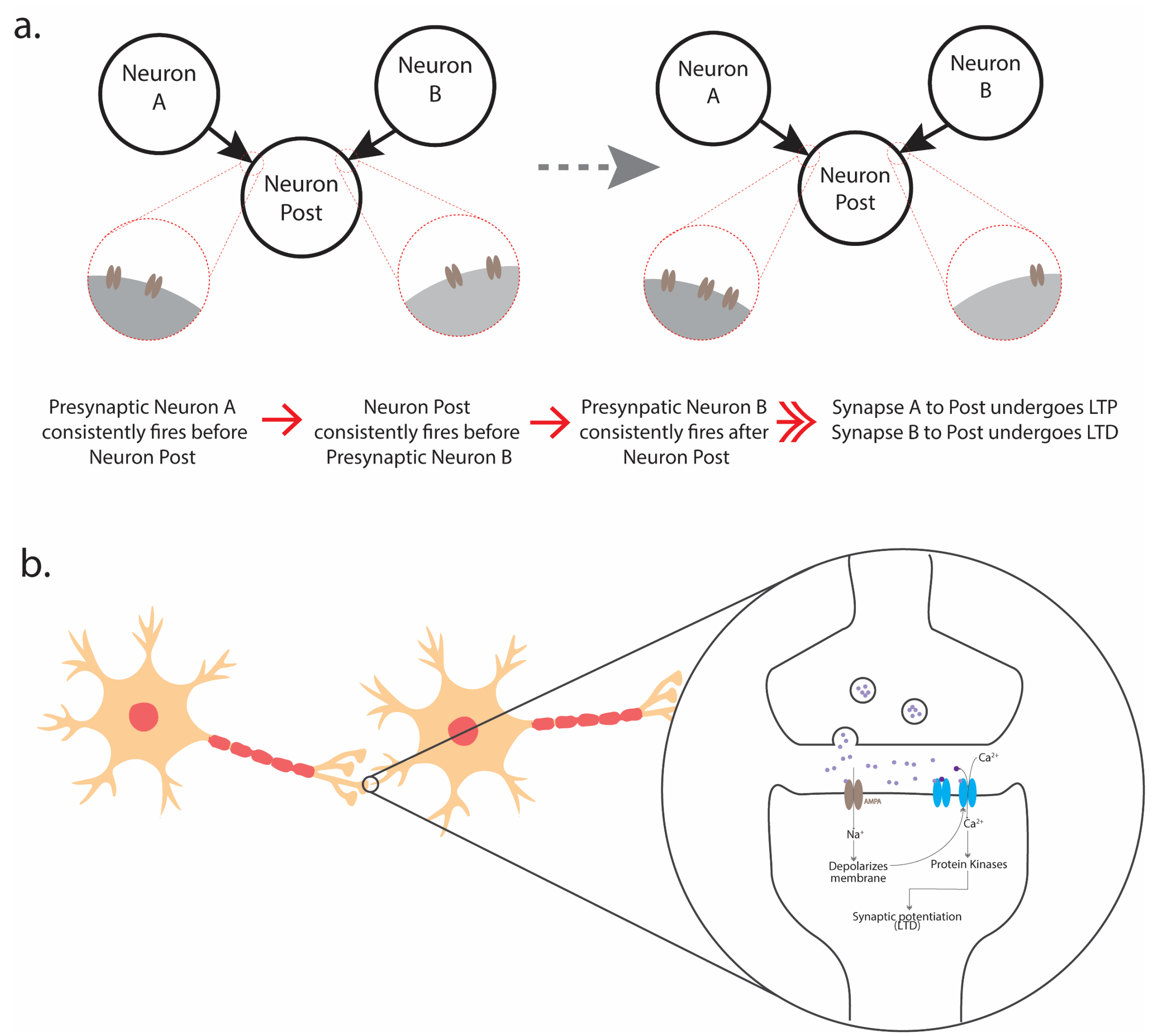


**Supplementary Figure 1**. **a** Representative diagram representing the induction of Spike Timing Dependent Plasticity **b** Representative diagram of AMPA/NMDA receptor mediated LTP

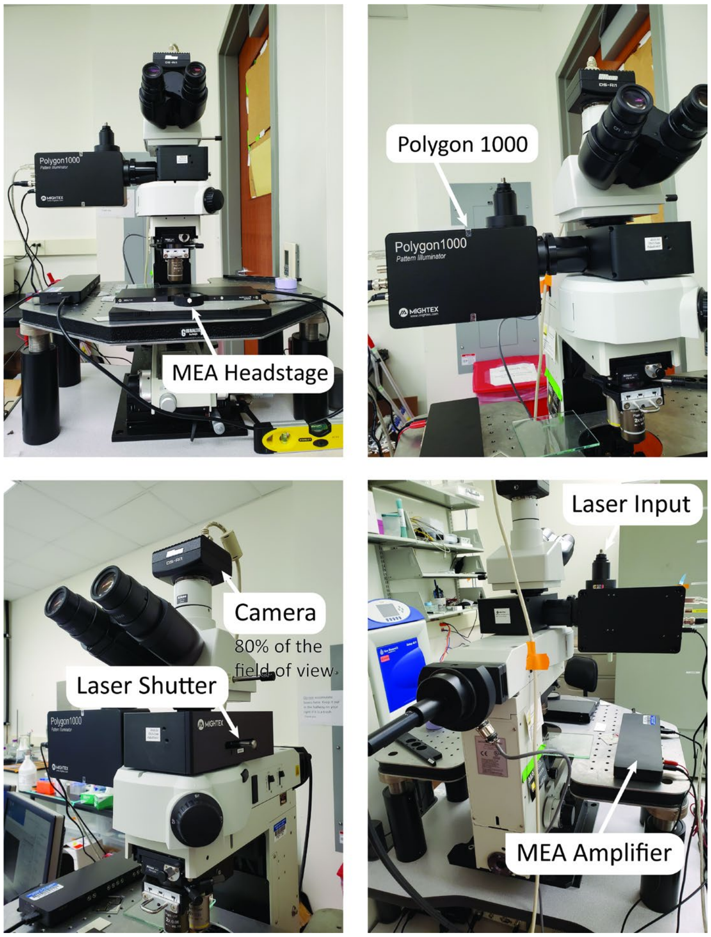


**Supplementary Figure 2**. Representative pictures of the microscope/stimulation system unto which the MEA recording system was attached to.

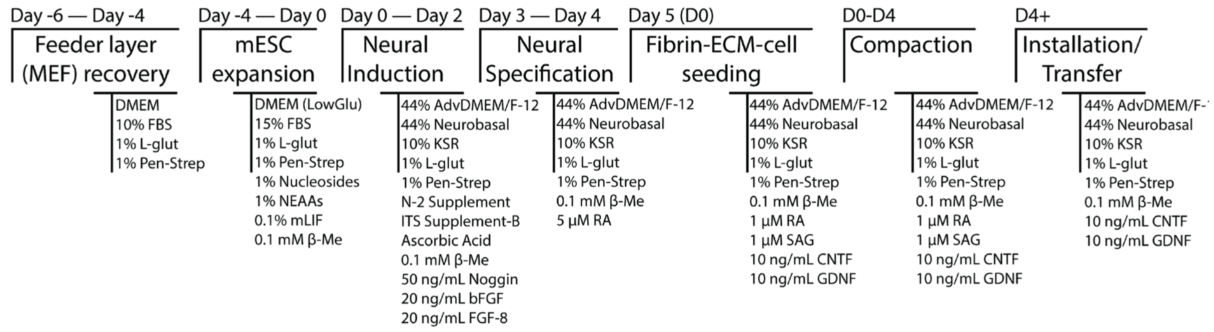


**Supplementary Figure 3. Differentiation protocol for increasing glutamatergic neuronal yield in mESC-derived EBs**

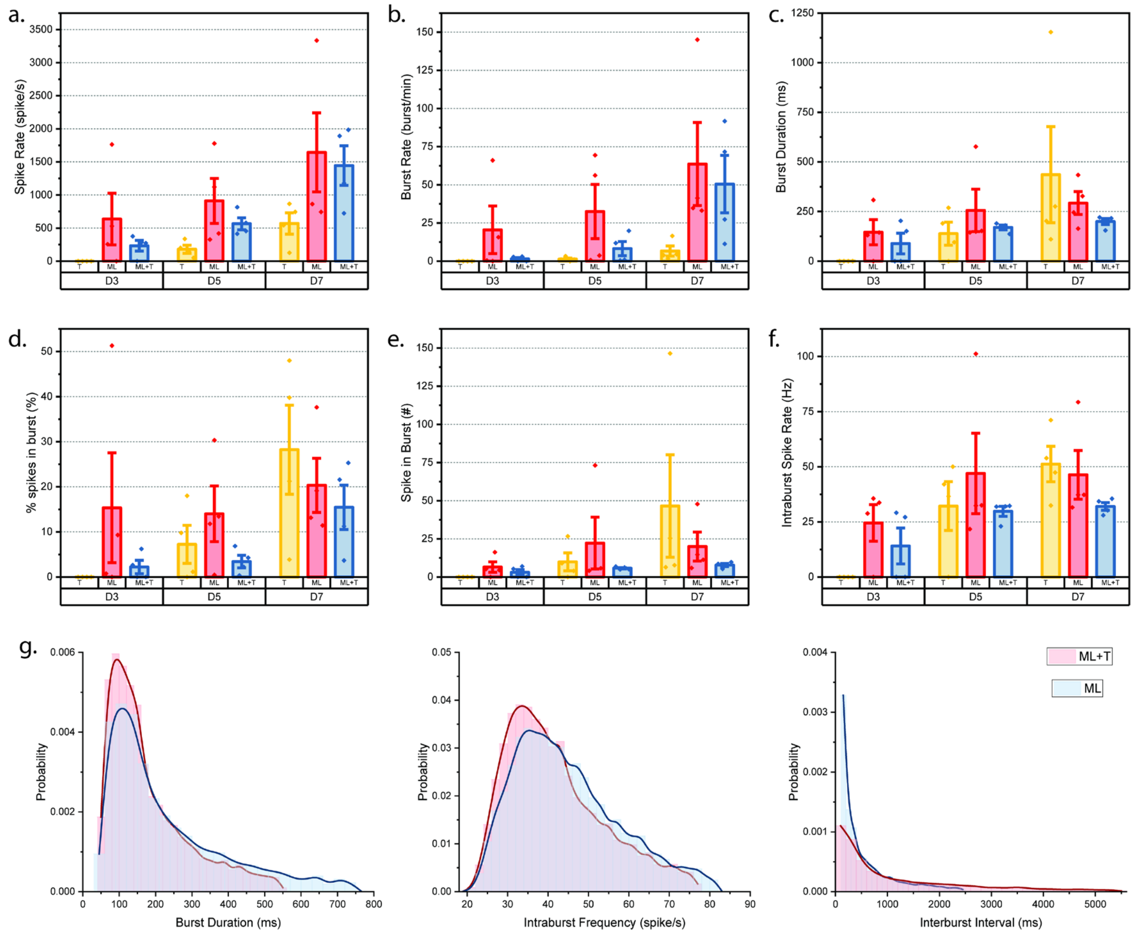


**Supplementary Figure 4. Burst parameters were compared between the monolayer of neurons (ML), NTM only (T) and NTM seeded on top of T (ML+T):** Rate of spiking **(a)**, rate of bursting **(b)**, the mean duration of bursts **(c)**, percentage of spikes that fire in bursts **(d)**, average number of spikes per burst **(e)** and average distance of spikes within bursts **(f)**. **g** PDFs of burst parameters between ML and ML+T recordings.

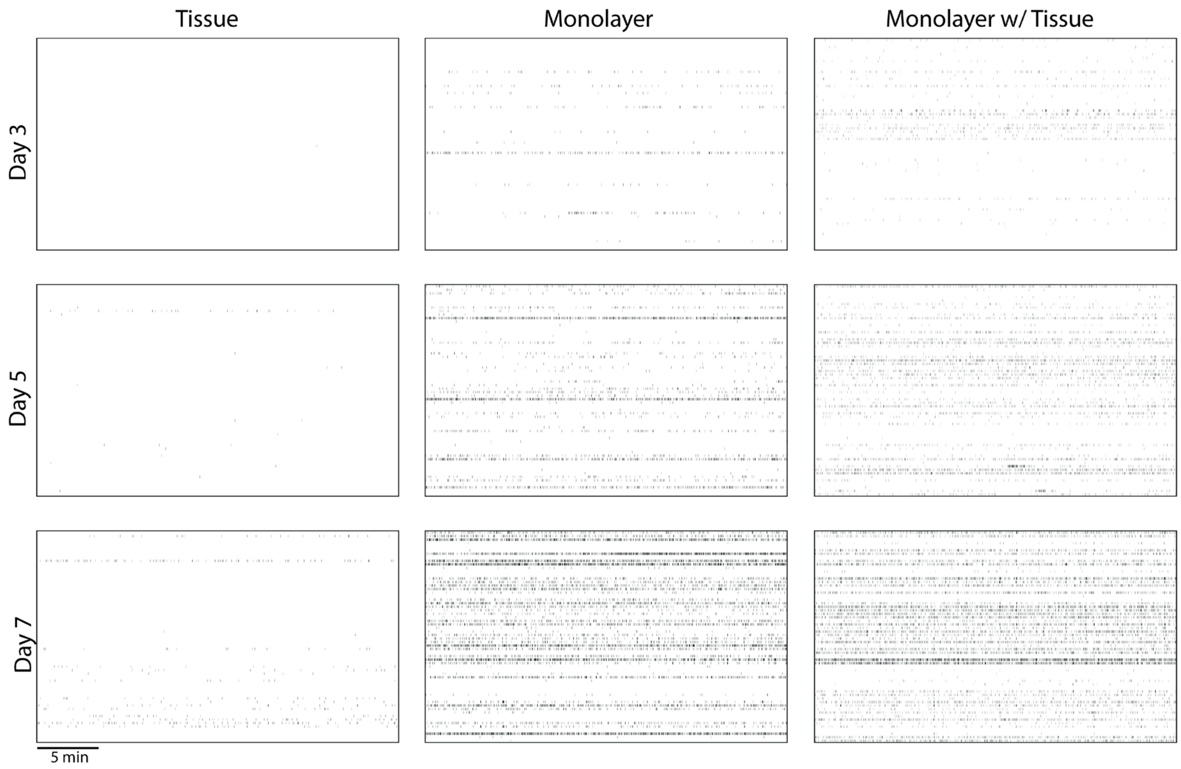


**Supplementary Figure 5. Representative raster plots of recordings between T, ML and ML+T**

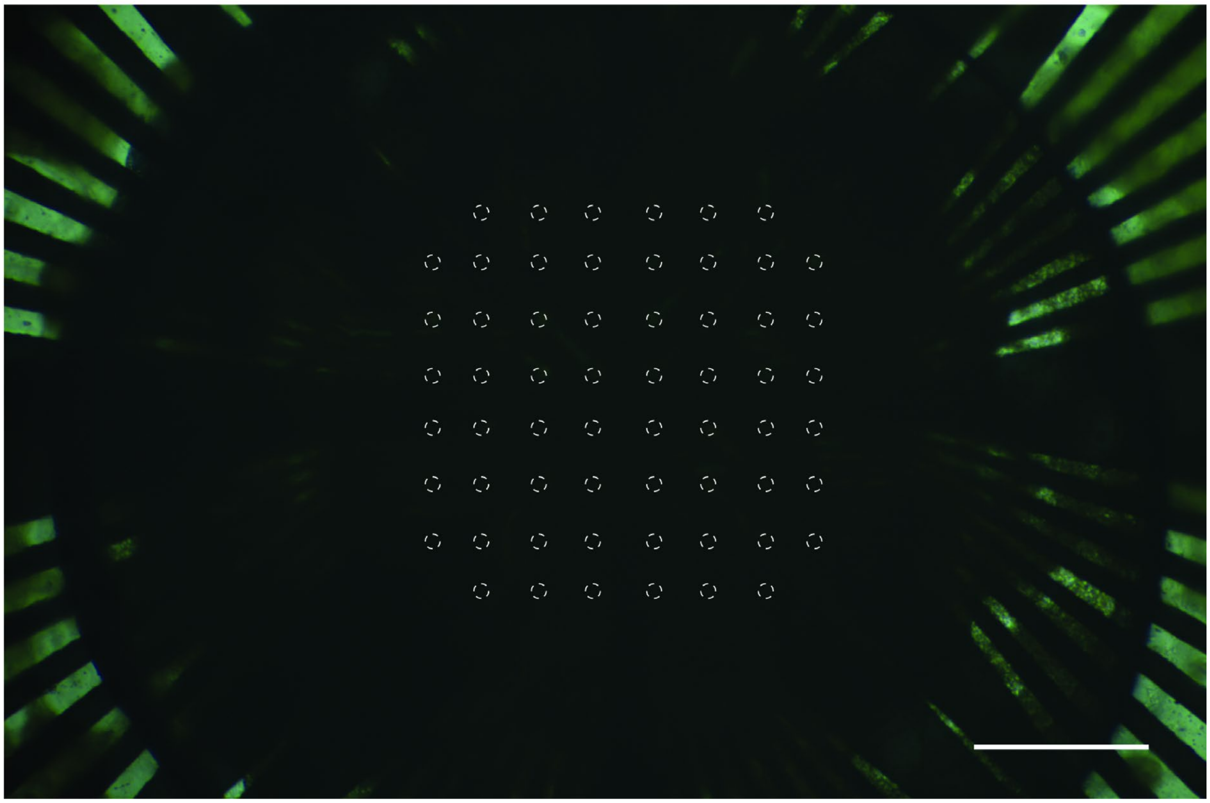

**Supplementary Figure 6. Representative picture of ML+T construct fabricated on-chip**

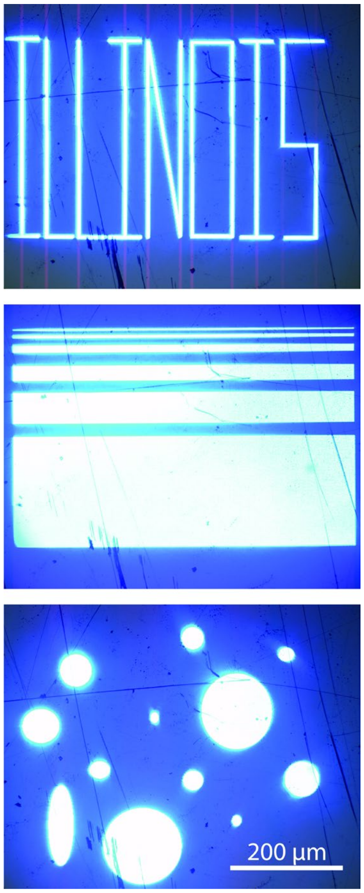


**Supplementary Figure 7. Representative examples showing the flexibility of stimulation design with Polygon System**

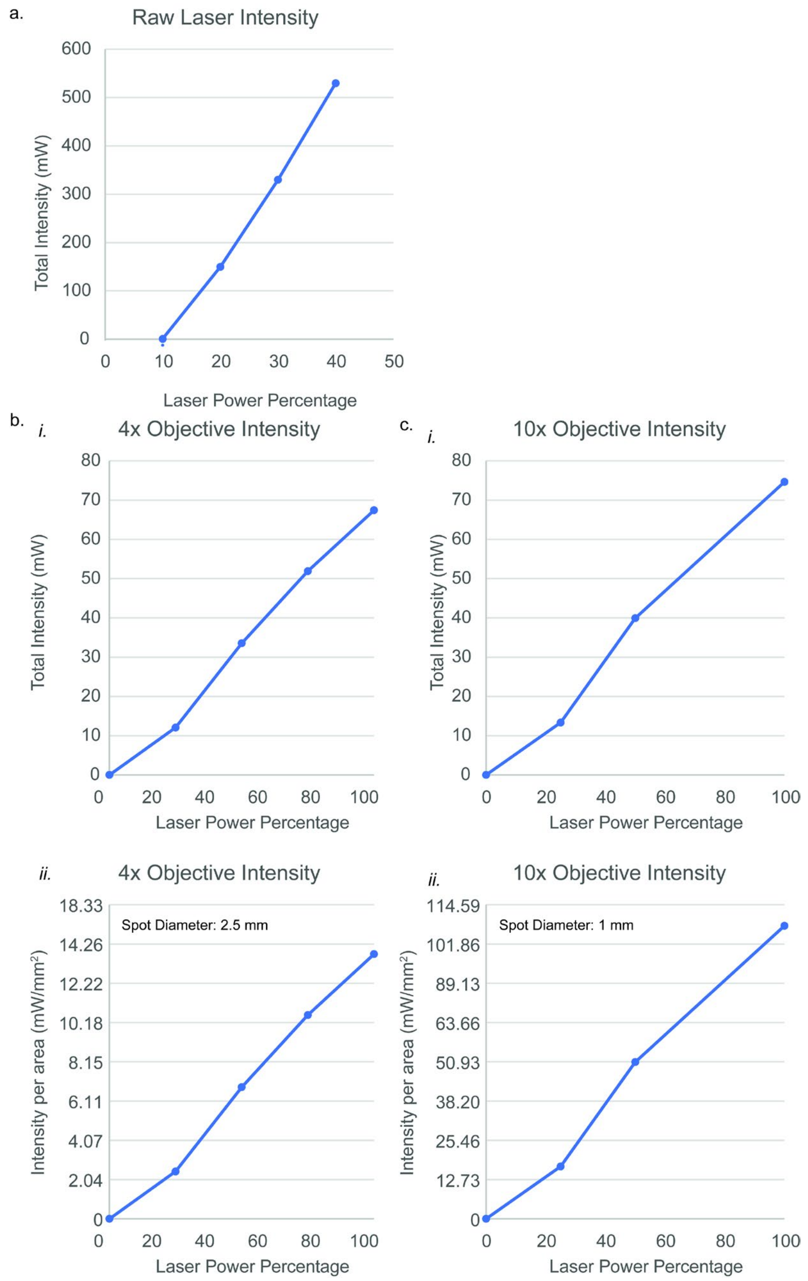


**Supplementary Figure 8. Polygon1000 power validation a** Resulting laser intensity of polygon at different laser powers prior to passing through microscope. **b** Resulting total laser intensity of polygon at different laser powers passing through 4X objective ***(i)*** as well as intensity per area (***ii***) **c** Resulting total laser intensity of polygon at different laser powers passing through 10X objective (***i)*** as well as intensity per area (***ii***)

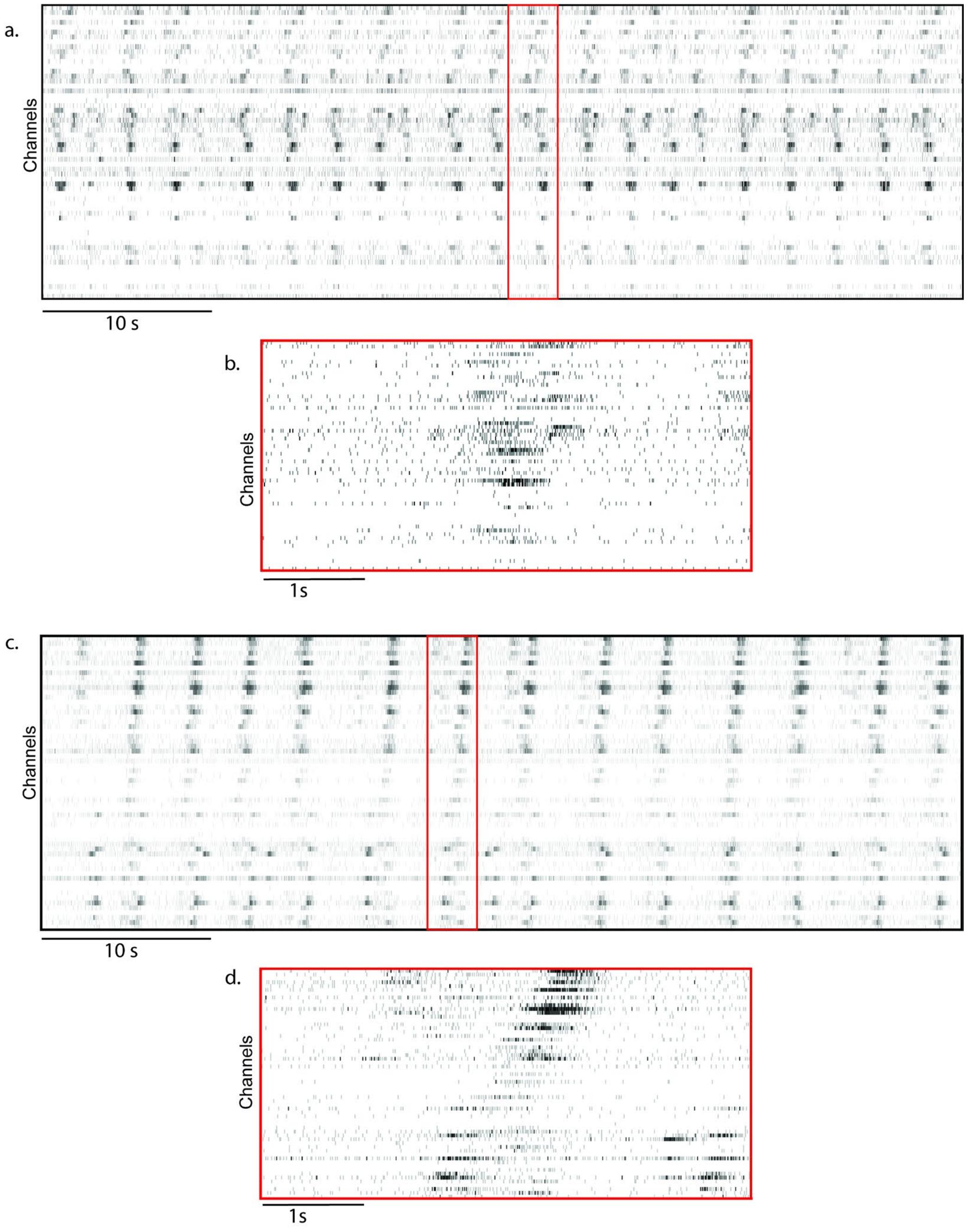


**Supplementary Figure 9. Representative raster plots of trained ML+T constructs on-chip a** 1 minute excerpt of spontaneous activity on the first day of training with a single bursting event magnified (**b**). **c** 1 minute excerpt of spontaneous activity on the third day of training with a single bursting event magnified (**d**).

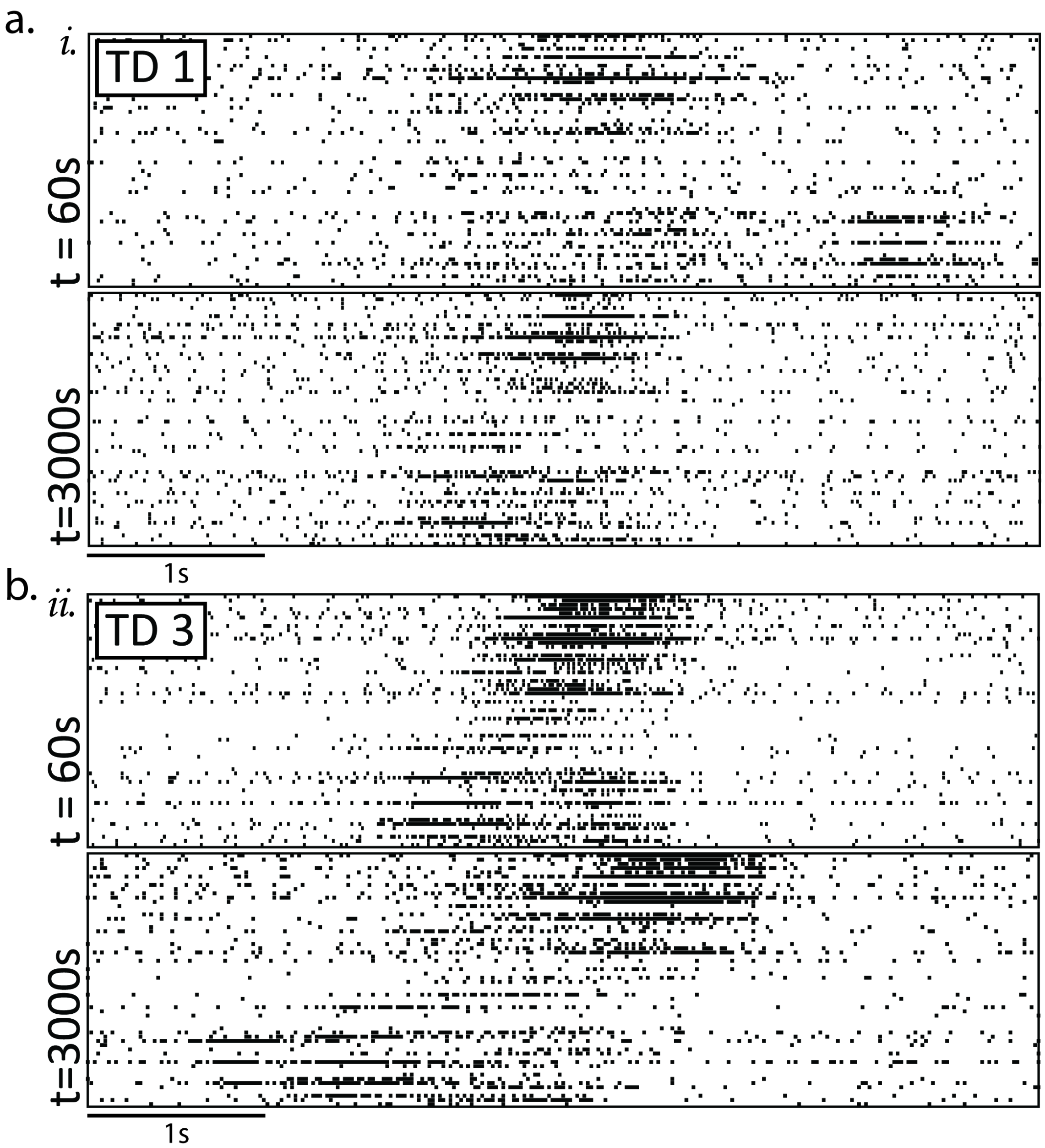


**Supplementary Figure 10. Resulting raster plots for training monolayer only of neurons. a** Raster plot of a representative burst on training day 1, 60 s into the recording before training (*i*) and at 3000s into the recording, at the end of training (*ii*). **b** Raster plot of a representative burst on training day 3, 60 s into the recording before training (*i*) and at 3000s into the recording, at the end of training (*ii*).
